# CB_1_R Regulates Soluble Leptin Receptor Levels via CHOP, Contributing to Hepatic Leptin Resistance

**DOI:** 10.1101/2020.07.14.202283

**Authors:** Adi Drori, Asaad Gammal, Shahar Azar, Liad Hinden, Rivka Hadar, Daniel Wesley, Alina Nemirovski, Gergő Szanda, Maayan Salton, Boaz Tirosh, Joseph Tam

**Author notes:** Address correspondence to: Joseph Tam, Obesity and Metabolism Laboratory, The Institute for Drug Research, School of Pharmacy, Faculty of Medicine, POB 12065, Jerusalem 91120, Israel.

## Abstract

The soluble isoform of leptin receptor (sOb-R), secreted by the liver, regulates leptin bioavailability and bioactivity. Its reduced levels in diet-induced obesity (DIO) contributes to hyperleptinemia and leptin resistance, effects that are known to be regulated by the endocannabinoid (eCB)/CB_1_R system. Here we show that pharmacological activation/blockade as well as genetic overexpression/deletion of hepatic CB_1_R modulates sOb-R levels and consequently hepatic leptin resistance. Interestingly, peripheral CB_1_R blockade failed to reverse DIO-induced reduction of sOb-R levels, fat mass, dyslipidemia, and hepatic steatosis in mice lacking C/EBP homologous protein (CHOP), whereas direct activation of CB_1_R in hepatocytes reduced sOb-R levels in a CHOP-dependent manner. Moreover, CHOP stimulation increased sOb-R expression and release via a direct regulation of its promoter, while CHOP deletion reduced leptin sensitivity. Our findings highlight a novel molecular aspect by which the hepatic eCB/CB_1_R system involves in the development of hepatic leptin resistance by regulating sOb-R levels via CHOP.

**Summary:** Here we describe a novel molecular aspect by which the hepatic endocannabinoid/CB_1_R system contributes to hepatic leptin resistance by regulating soluble leptin receptor levels via CHOP.

## Introduction

Leptin, predominantly produced by and secreted from white adipocytes, conveys information regarding the status of energy storage and availability to the brain, to maintain energy homeostasis. It binds the leptin receptor in hypothalamic neurons to reduce food intake and increase energy expenditure in coordination with other adipokines and gastric peptides (Allison & Myers, 2014; Pan & Myers, 2018). Molecularly, leptin stimulates the secretion of α-melanocortin stimulating hormone (α-MSH) from proopiomelanocortin (POMC) neurons at the arcuate nucleus (ARC), and inhibits the secretion of the orexigenic peptides neuropeptide-Y (NPY) and Agouti-related protein (AgRP) (Flak & Myers, 2016). Genetic leptin deficiency or lack of functional leptin receptor results in morbid obese and insulin resistance phenotypes in animals (*ob/ob* or *db/db* mice, respectively) (Tartaglia et al., 1995; Y. Zhang et al., 1994). In humans, congenital leptin deficiencies are rare, leading to hyperphagia and early-onset obesity, which can be reversed with a leptin replacement therapy (Mantzoros, 1999). However, most cases of obesity are characterized by hyperleptinemia, indicating that obesity is a leptin-resistant state, where leptin signaling is impaired.

Various mechanisms have been linked to the development of diet-induced obesity (DIO)-related leptin resistance, including limited CNS access of leptin due to saturated transport machinery, uncoupling of leptin from its receptor (due to rare genetic mutations or intra-cellular modulators), leptin-induced downregulation of its hypothalamic receptor, and several circulating factors such as the soluble isoform of leptin receptor (sOb-R) [reviewed in (Engin, 2017; Martin, Qasim, & Reilly, 2008)]. Both in humans and mice, the leptin receptor gene (*LEPR*) encodes 4 membrane-anchored isoforms, which differ in the length of their cytoplasmic tail. The long isoform, Ob-Rb, is considered to convey the most robust cellular response to leptin, while the shorter isoforms (Ob-Ra, Ob-Rc, and Ob-Rd) carry a weaker signal. In addition, sOb-R, which lacks the trans-membrane and intra-cellular domains, also exists. In humans, sOb-R is exclusively generated via proteolytic shedding of membrane-anchored isoforms (Maamra et al., 2001), whereas in mice, it is produced by both transcription of a designated isoform (Ob-Re) and ectodomain shedding of Ob-Rb and Ob-Ra (Ge, Huang, Pourbahrami, & Li, 2002; Li, Ioffe, Fidahusein, Connolly, & Friedman, 1998). sOb-R, mainly produced by hepatocytes, is the main leptin-binding protein in human plasma, regulating leptin’s bioavailability and bioactivity (Lammert, Kiess, Bottner, Glasow, & Kratzsch, 2001; G. Yang, Ge, Boucher, Yu, & Li, 2004). In fact, studies have shown that the circulating levels of sOb-R are inversely correlated with body weight and free leptin levels (Ogier, Ziegler, Mejean, Nicolas, & Stricker-Krongrad, 2002). In addition, sOb-R levels are increased following weight loss (Laimer et al., 2002; Reinehr, Kratzsch, Kiess, & Andler, 2005), and its overexpression in mice increases leptin sensitivity (Huang, Wang, & Li, 2001; Lou et al., 2010), supporting the key role of sOb-R in the development as well as the reversal of leptin resistance.

The endocannabinoid (eCB) system, a major regulator of energy homeostasis (Fride, Bregman, & Kirkham, 2005; Pagotto, Marsicano, Cota, Lutz, & Pasquali, 2006), evokes various cellular/metabolic pathways via the activation of two G-protein-coupled receptors, cannabinoid type-1 (CB_1_R) and type-2 (CB_2_R) receptors, by the main eCBs, *N*-arachidonoylethanolamine (AEA) and 2-arachidonoylglycerol (2-AG). The eCB/CB_1_R system is highly overactive during obesity (Engeli, 2008; Engeli et al., 2005), and both central and peripheral stimulations of this system have been suggested to contribute to the development of the metabolic syndrome, including leptin resistance (Engeli et al., 2005; Pagotto, Vicennati, & Pasquali, 2005). Studies have shown that leptin’s ability to regulate food intake and peripheral lipid metabolism depends upon hypothalamic CB_1_Rs (Buettner et al., 2008; Cardinal et al., 2014; Di Marzo et al., 2001; Jo, Chen, Chua, Talmage, & Role, 2005; Malcher-Lopes et al., 2006). Recent evidence demonstrates that peripheral CB_1_R signalling has the ability to modulate leptin activity too. By using peripherally restricted CB_1_R blockers, we have recently demonstrated that DIO-related hyperleptinemia is completely reversed by increasing leptin’s renal clearance and decreasing its secretion from adipocytes (Tam et al., 2012; Tam et al., 2010). Additionally, we have shown that the reversal of hypothalamic leptin resistance in obese mice treated with the peripherally restricted CB_1_R blocker, JD5037, is mediated via re-sensitizing the animals to endogenous leptin and re-activating POMC neurons (Tam et al., 2017). However, a direct role for the eCB/CB_1_R system in the regulation of sOb-R levels and hepatic leptin signaling has never been investigated.

## Results

### Hepatic CB_1_R regulates sOb-R levels and leptin signalling

To evaluate the direct contribution of CB_1_R to the regulation of sOb-R levels, we first utilized a pharmacological inhibition paradigm of CB_1_R in DIO mice by using the peripherally restricted CB_1_R inverse agonist JD5037. Similarly to previous findings (Mazor et al., 2018), a significant reduction in serum levels of sOb-R was documented in obese mice; an effect that was ameliorated by JD5037 treatment (Figure 1A). Since sOb-R is mainly produced by the liver (Lammert et al., 2001), we also analyzed the content of sOb-R in liver homogenates from these animals and found a similar trend as in serum (Figure 1B). qPCR measurements of the *Ob-Re* isoform revealed that JD5037 treatment also affected its transcription and protein levels (Figure 1C-E). Moreover, the protein expression of two additional isoforms of LEPR (Ob-Rb and Ob-Ra) in liver homogenates was also decreased in DIO mice, and normalized following JD5037 treatment (Figure 1D-E).

**Figure 1.**
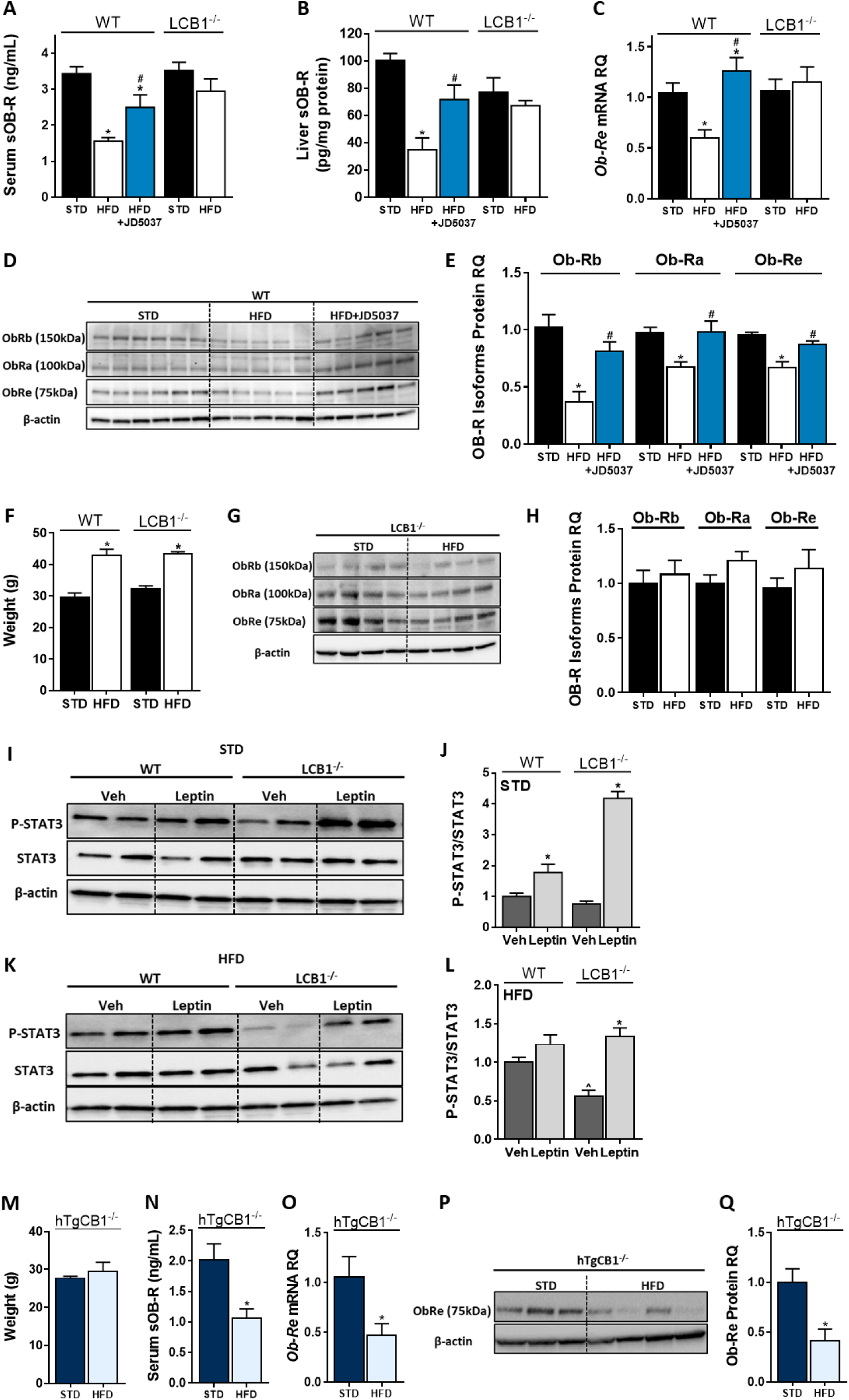
Hepatic CB_1_R regulates sOb-R levels and leptin signaling in DIO. Serum. (A**, n=10-**21) and liver **(B**, n=7-10) levels of sOb-R are reduced following a 14-weeks consumption of HFD in WT, but not LCB1^-/-^ mice. JD5037 (3 mg/kg, for 7 days) reverses the reduction in WT mice. The same trend observed in hepatic mRNA levels of *Ob-Re* **(C**, n=10-12) as well as protein level of Ob-Rb, Ob-Ra and Ob-Re **(D-E**, n=6-10; **G-H**, n=8-10). Despite a comparable weight gain following HFD consumption in WT and LCB_1_^-/-^ mice **(F**, n=12-18), obese mice that lack CB_1_R in hepatocytes remain leptin sensitive indicated by increased pSTAT3 levels **(I-L**, n=5). Transgenic mice, expressing CB_1_R only in hepatocytes, are protected from DIO **(M**, n=4-5). An exclusive hepatic expression of CB_1_R is sufficient for HFD feeding to induce reduction in serum, liver mRNA and protein levels of sOb-R **(N-Q**, n=4-5). Data represent mean ± SEM of indicated number of replicates in each panel. *P<0.05 relative to STD fed animals from the same strain. #P<0.05 relative to HFD fed mice from the same strain. ^P<0.05 relative to the same treatment group of WT mice.

To further establish the contribution of hepatic CB_1_R to the high-fat diet (HFD)-induced decrease in sOb-R levels, we utilized the liver-specific CB_1_R null (LCB1^-/-^) mice, a genetic deletion model of mice that lack CB_1_R specifically in hepatocytes [mouse model generation is described in (Osei-Hyiaman et al., 2008)]. When fed with a HFD, these mice gain similar weight to their wild-type (WT) littermate controls [(Osei-Hyiaman et al., 2008) and Figure 1F]; however, they are less prone to develop liver steatosis, dyslipidemia and leptin resistance (Osei-Hyiaman et al., 2008), making hepatic CB_1_R a central regulator of obesity-related liver complications. We were therefore not surprised to find that the liver specific deletion of CB_1_R was sufficient to maintain normal circulating levels of sOb-R in both lean and obese LCB1^-/-^ mice (Figure 1A). Similarly, the hepatic gene and protein expression of sOb-R and the other LEPR isoforms were not affected by the HFD feeding (Figure 1B, C and G, H), suggesting that hepatic CB_1_R most likely regulates sOb-R levels.

To test the functional relevance of our findings to hepatic leptin signalling, we measured the phosphorylation levels of STAT3, the gold-standard measure of leptin signalling [reviewed in (Allison & Myers, 2014)], in mouse livers following exogenous leptin administration *in vivo*. Whereas both lean WT and LCB1^-/-^ mice showed elevated pSTAT3/STAT3 ratio in response to leptin (Figure 1I, J), only obese LCB1^-/-^ mice remained leptin sensitive (Figure 1K, L). These results are in line with findings from Osei-Hyiaman and colleagues (Osei-Hyiaman et al., 2008), demonstrating that LCB1^-/-^ mice are resistant to obesity-induced hyperleptinemia.

Additional support for the regulation of sOb-R by hepatic CB_1_R derived from another transgenic mouse model (hTgCB1^-/-^), in which CB_1_R is expressed only in hepatocytes [mouse model generation is described in (Liu et al., 2012; Tam et al., 2010)]. These mice, despite being resistance to DIO like global CB_1_R^-/-^ mice [(Liu et al., 2012; Tam et al., 2010) and Figure 1M], demonstrate increased circulating leptin levels when fed a HFD (Liu et al., 2012). In accordance with that, the circulating and hepatic sOb-R levels in these mice were decreased by 50% following 14 weeks consumption of a HFD (Figure 1N-Q). Hence, over expression of CB_1_R in the liver alone recapitulates the HFD-induced downregulation of sOb-R observed in WT mice.

Next, we assessed whether a direct activation of CB_1_R in hepatocytes induces a reduction in sOb-R levels. To test this, we treated cultured hepatocytes with the synthetic CB_1_R agonist noladin ether (NE) for 24 hrs. We analysed both culture media and cell lysates, and found that, similarly to obesity, direct activation of CB_1_R also decreased sOb-R levels in the culture media. This was also the case with intra-cellular levels of other LEPR isoforms measured. This CB_1_R-mediated reduction in ObR levels was completely reversed by blocking CB_1_R using JD5037 (Figure 2).

**Figure 2.**
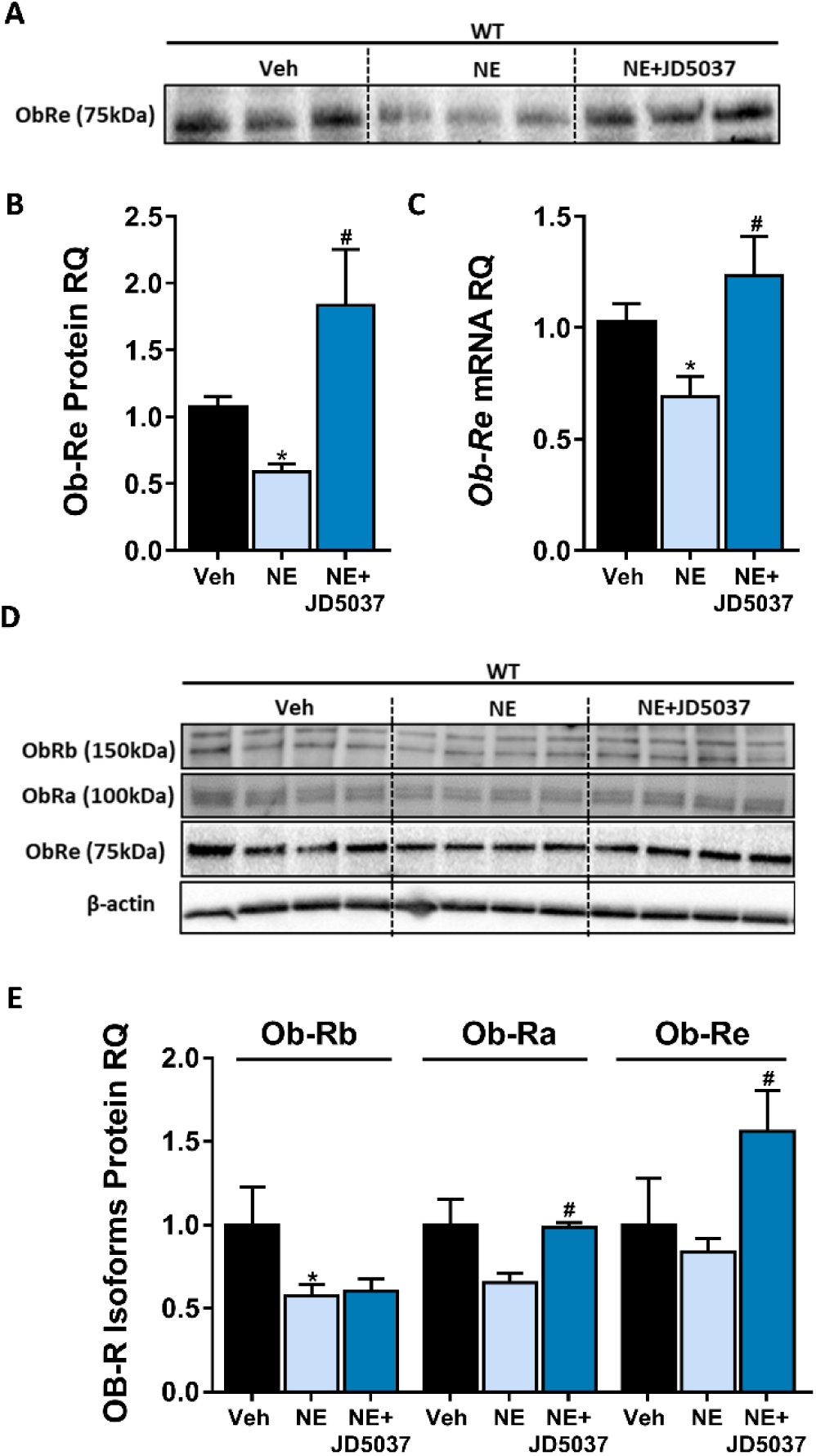
CB_1_R directly regulates sOb-R levels in hepatocytes. 24h treatment with synthetic CB_1_R agonist noladin ether (NE, 2.5 μM) induced reduction in sOb-R levels in the culture media of immortalized hepatocytes (blot was quantified using Ponceau staining as a loading control). This was completely ameliorated by 1 hr pre-treatment with 100 nM JD5037 **(A-B)**. Similar results were observed in both mRNA **(C)** and protein **(D-E)** levels in hepatocytes lysate (for ObRb, the lower band was quantified). Data represent mean ± SEM of 12-15 replicates from at least 3 independent experiments. Blots are representative. *P<0.05 relative to vehicle treated cells. #P<0.05 relative to NE-treated cells.

### CHOP contributes to the metabolic response to peripheral CB_1_R blockade

Several lines of evidence suggest that hypothalamic neurons, including POMC, experience endoplasmic reticulum (ER) stress during DIO, which may contribute to the development of leptin resistance (L. Ozcan et al., 2009; Ramirez & Claret, 2015). We have previously reported that pharmacological inhibition of peripheral CB_1_Rs (by AM6545) reverses the HFD-induced hepatic elevation in the ER stress marker phospho-eIF2α (Tam et al., 2010). Since ER stress strongly affects protein translation and secretion [reviewed in (Ron & Walter, 2007)], we hypothesized that it could explain the reduced expression/secretion of sOb-R following CB_1_R activation and/or obesity. We therefore measured the phosphorylation of eIF2α and found the expected elevation in phospho-to-total eIF2α ratio in the liver of HFD fed WT mice. Similar to AM6545, treatment with JD5037 normalized p-eIF2α levels (Supplementary Figure 1A, B), suggesting relived ER stress following CB_1_R blockade. In agreement with these findings, a comparable ratio of hepatic phospho-to-total eIF2α ratio was documented in lean and obese LCB1^-/-^ mice (Supplementary Figure 1C, D).

Measuring the expression levels of the ER stress marker C/EBP homologous protein (CHOP) revealed surprising findings, since both the hepatic mRNA and protein levels of CHOP were downregulated in obese WT mice, despite the suggested ER stress. Its expression levels were even further upregulated by JD5037, and remained comparable between lean and obese LCB1^-/-^ mice (Figure 3A-D). In fact, CHOP levels were positively correlated with the levels of sOb-R in both our experimental paradigms, leading us to hypothesize that CHOP may directly be involved in the regulation of sOb-R.

**Figure 3.**
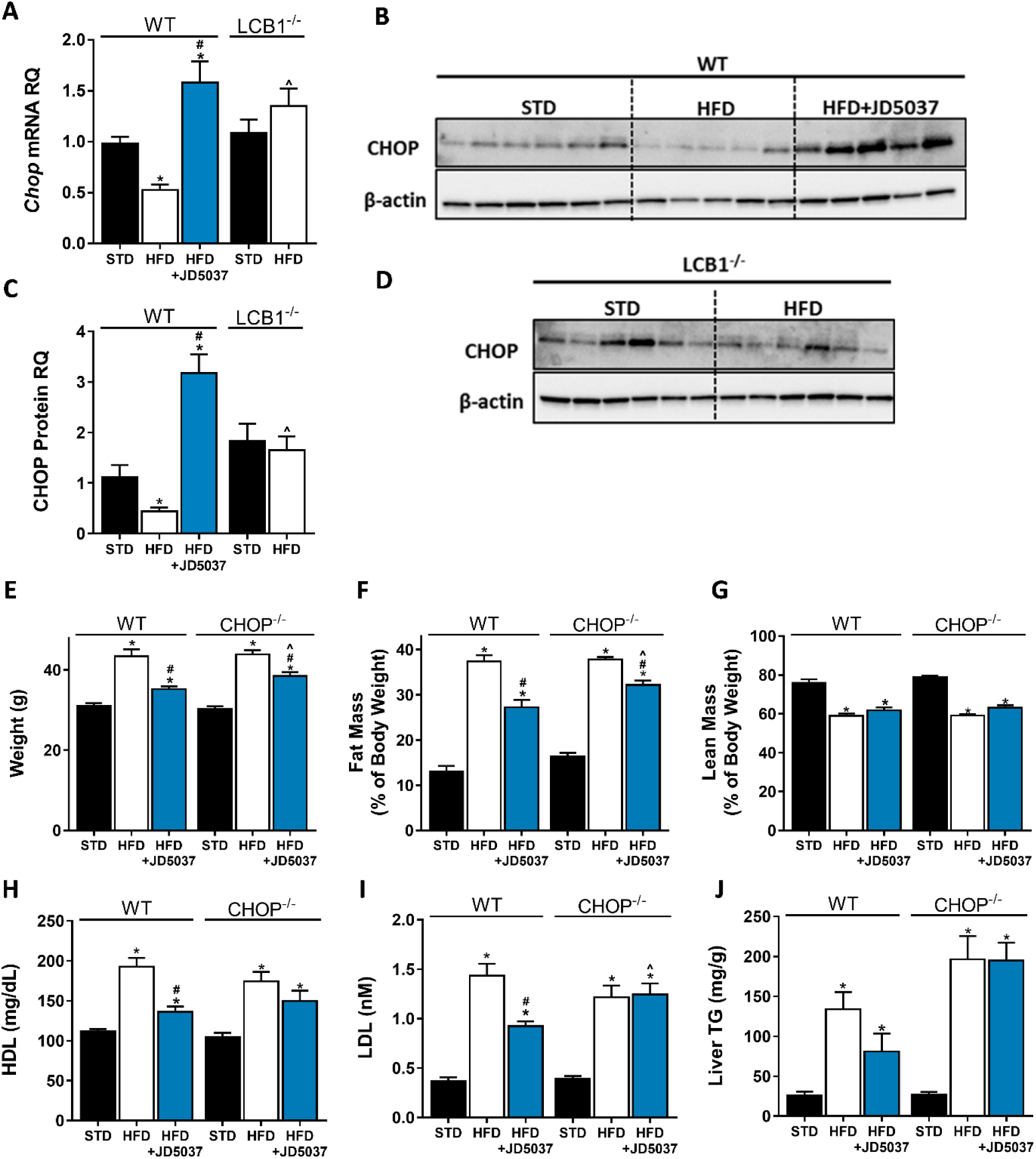
CHOP contributes to the metabolic benefits of peripheral CB_1_R blockade. mRNA **(A**, n=8-17) and protein **(B-D**, n=5-6) levels of CHOP show reduced expression following 14 weeks on HFD in WT, but not LCB1^-/-^ mice. JD5037 (3 mg/kg, for 7 days) treatment reverses the HFD-induced reduction in CHOP levels. Metabolic assessment of mice revealed diminished effect of JD5037 in CHOP KO mice. Weight **(E**, n=15-20), fat and lean mass **(F & G**, n=8-20), serum HDL, LDL as well as hepatic triglycerides (TG) **(H-J**, n=15-20) were comparable in lean and obese WT and CHOP KO mice. JD5037 treatment was significantly more efficient in reducing weight, fat mass, LDL and TG in WT mice. Data represent mean ± SEM of indicated number of replicates in each panel. *P<0.05 relative to STD fed animals from the same strain. #P<0.05 relative to HFD fed mice from the same strain. ^P<0.05 relative to the same treatment group of WT mice.

To test our hypothesis, we compared the metabolic efficacy of JD5037 in obese CHOP KO mice and their littermate controls. Whereas JD5037 was almost equieffective in reducing body weight and fat mass in both obese mouse strains (Figure 3E-G), it improved plasma cholesterol levels as well as hepatic steatosis in WT mice only (Figure 3H-J). The reduced ability of peripherally restricted CB_1_R blockade to improve dyslipidemia and hepatic steatosis in CHOP KO mice led us to measure the hepatic eCB ‘tone’ in these mice. Strikingly, we found that the basal levels of AEA and 2-AG were markedly higher in CHOP KO mice than in the WT control group. Moreover, the increased eCB levels in CHOP KO mice remained unchanged following a consumption of HFD as well as JD5037 treatment (Supplementary Figure 2A, B). This could be partially explained by the differences documented in the mRNA expression patterns of fatty acid amide hydrolase (FAAH), monoacylglycerol lipase (MGLL), *N*-acyl phosphatidylethanolamine phospholipase D (NAPE-PLD), and diacylglycerol lipase alpha (DAGLα), the degrading and synthesizing enzymes of both eCBs, respectively (Supplementary Figure 2C-F). Overall, these data indicate that CHOP deficiency alters eCB ‘tone’, which, in turn, may lead to the reduced response to CB_1_R blockade mainly in the liver.

### CHOP plays a key role in the regulation of sOB-R by the eCB/CB_1_R system

Measuring the effect of CHOP deficiency on sOb-R levels revealed comparable circulating levels of sOB-R in lean and obese mice in the two mouse strains. However, JD5037 failed to restore sOb-R levels in CHOP KO mice (Figure 4A). The assessment of sOb-R mRNA expression and protein content in the livers of both strains documented reduced baseline levels in CHOP KO mice, compare to WT, which still remained low following HFD consumption and/or JD5037 treatment (Figure 4B, C). A similar trend was observed in the protein level of two more LEPR isoforms (Figure 4D, E). The HFD-induced hyperleptinemia was almost completely normalized by JD5037 treatment in WT mice, whereas it was only partially ameliorated by JD5037 in CHOP KO animals (Figure 4F). Taken together, our data suggest that regulation of ObR levels is CHOP-dependent. In addition, CHOP regulation of the soluble isoform levels can consequently affect circulating leptin levels, possibly, in a CB_1_R-dependent manner.

**Figure 4.**
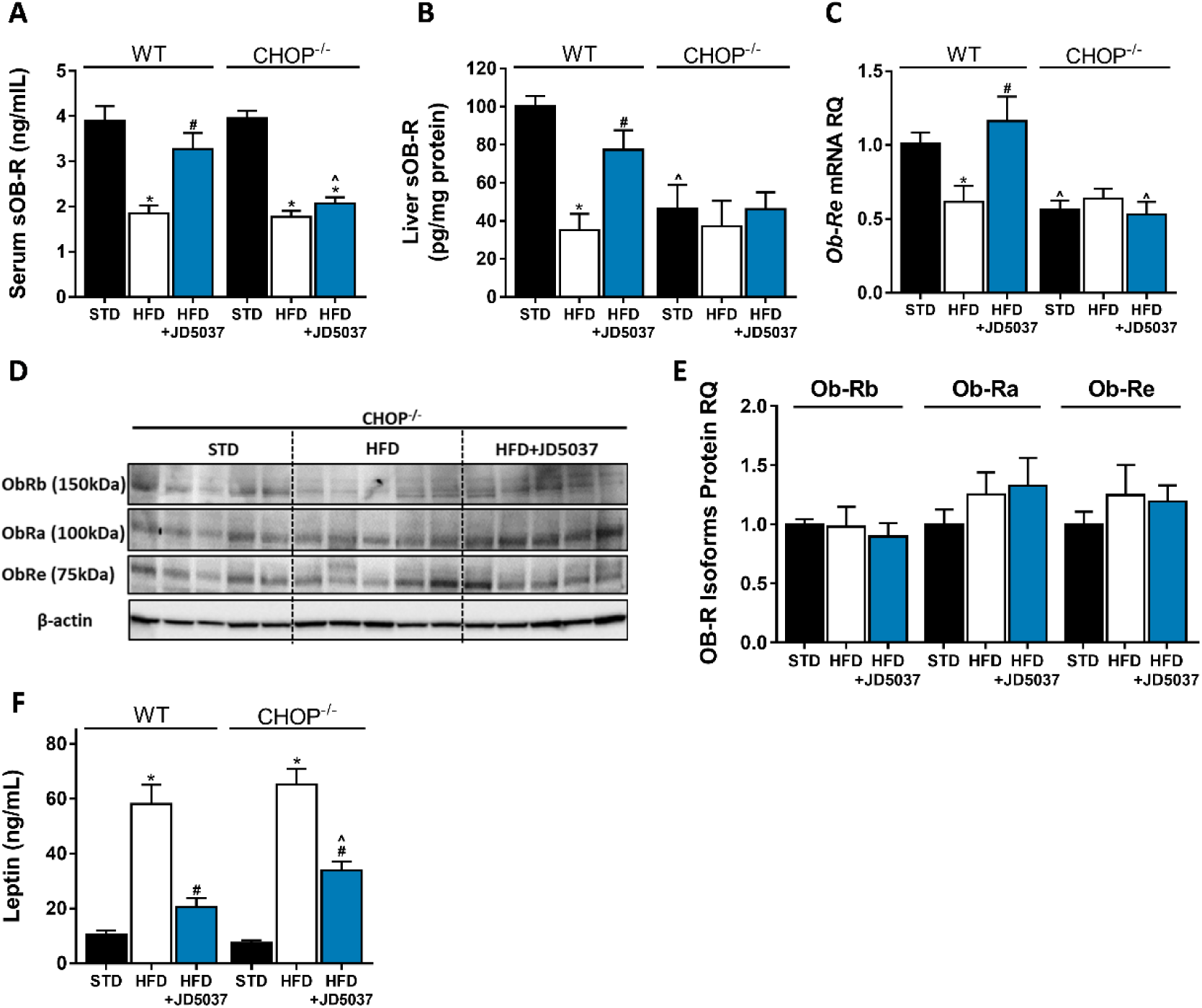
CHOP plays a key role in regulating sOB-R by the eCB/CB_1_R System. Serum **(A**, n=9-15) levels of sOb-R were reduced following a 14-weeks consumption of HFD. JD5037 (3 mg/kg, for 7 days) reversed the reduction in WT, but not CHOP KO mice. Basal hepatic levels of sOb-R were lower in CHOP KO and did not change following HFD or JD5037 treatment **(B**, n=7-10). A similar trend was observed in hepatic mRNA levels **(C**, n=9-16) and protein level of Ob-Rb, Ob-Ra and Ob-Re **(D & E**, n=5-6). Whereas DIO-related hyperleptinemia was comparable between WT and CHOP KO mice, JD5037 was more efficacious in reducing it in WT mice **(F**, n=9-16). Western blots are representative. Data represent mean ± SEM of indicated number of replicates in each panel. *P<0.05 relative to STD fed animals from the same strain. #P<0.05 relative to HFD fed mice from the same strain. ^P<0.05 relative to the same treatment group of WT mice.

To further investigate this concept, we directly activated CB_1_R (with NE) in immortalized hepatocytes originated from WT or CHOP KO mice. Similar to a HFD consumption in mice (Figure 3A), a direct activation of CB_1_R downregulated CHOP mRNA expression (Figure 5A). We validated this by measuring the expression levels of *Gadd34*, a downstream target of CHOP (Hu, Tian, Ding, & Yu, 2018), and found that its expression was also reduced in NE-treated WT mice, and remained unchanged in CHOP KO cells (Figure 5B), suggesting that CB_1_R activation in fact leads to reduced CHOP expression and activity. Whereas NE was able to reduce sOb-R levels in WT hepatocytes, it had the opposite effect in CHOP KO hepatocytes, suggesting that CB_1_R may regulate sObR levels in other mechanisms independently of CHOP (Figure 5C-E).

**Figure 5.**
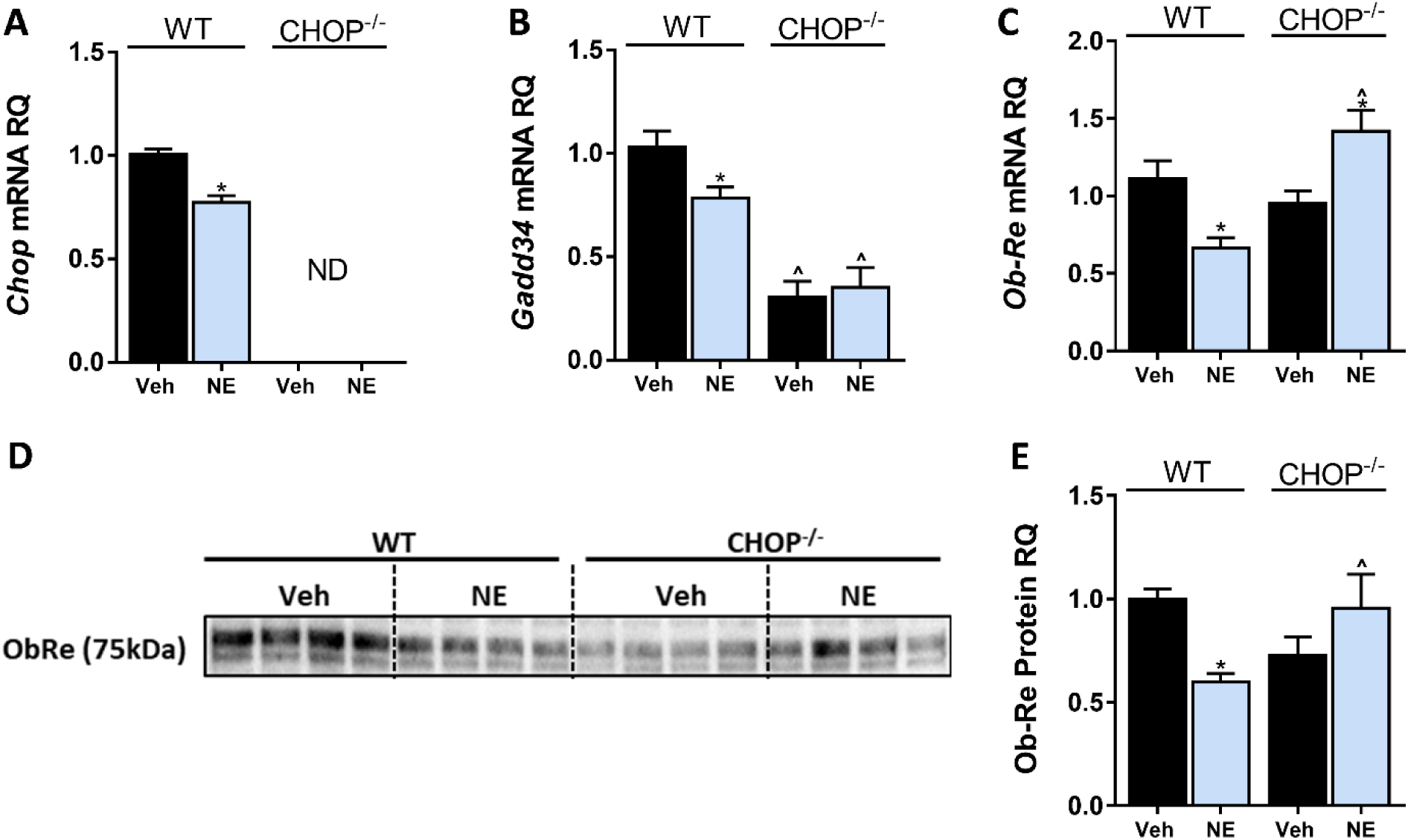
CHOP regulates LEPR expression in hepatocytes. *In vitro*, 24h treatment with noladin ether (NE; 2.5 μM) induced reduction in mRNA levels of *Chop.* <u>(ND-Not Detected)</u> **(A)**, *Gadd34* **(B)**, and *Ob-Re* **(C)** in WT, but not CHOP KO hepatocytes. A similar trend was observed in sOb-R protein levels secreted into the culture media of hepatocytes (blot was quantified using Ponceau staining as a loading control) **(D, E)**. Data represent mean ± SEM of 12-15 replicates from at least 3 independent experiments. Blots are representative. *P<0.05 relative to vehicle treated cells in the same genotype. ^P<0.05 relative to same treatment paradigm in WT.

The consistent correlation between CHOP and sOb-R levels implies that CHOP is a positive regulator of Ob-Re. To validate this further, we analysed Ob-Re levels in WT and CHOP KO hepatocytes treated with tunicamycin (TM), a potent inducer of ER stress. Treatment with TM for 6 hrs led to an expected and robust expression of CHOP mRNA and protein in WT cells (Figure 6A, B). Importantly, this was accompanied with elevated mRNA expression levels of Ob-Re as well as secreted levels of sOb-R into the culture media in WT, but not CHOP KO hepatocytes (Figure 6C-E). Increased levels of sOb-R in culture media were also documented when we exogenously overexpressed myc-tagged CHOP in WT hepatocytes (Figure 6F, G), supporting a direct role for CHOP in LEPR gene regulation. By using a luciferase reporter assay, in which the −650 to +850 (relative to transcription start site) region of the LEPR promoter was cloned into firefly luciferase expressing vector, we found that CHOP expression and luciferase activity in transfected cells was induced using TM (Figure 6H), while CB_1_R activation using NE (which downregulates CHOP expression as seen in Figure 5A) had an opposite effect in WT, but not in CHOP KO cells. These data support the involvement of CHOP in CB_1_R-dependent regulation of sOb-R.

**Figure 6.**
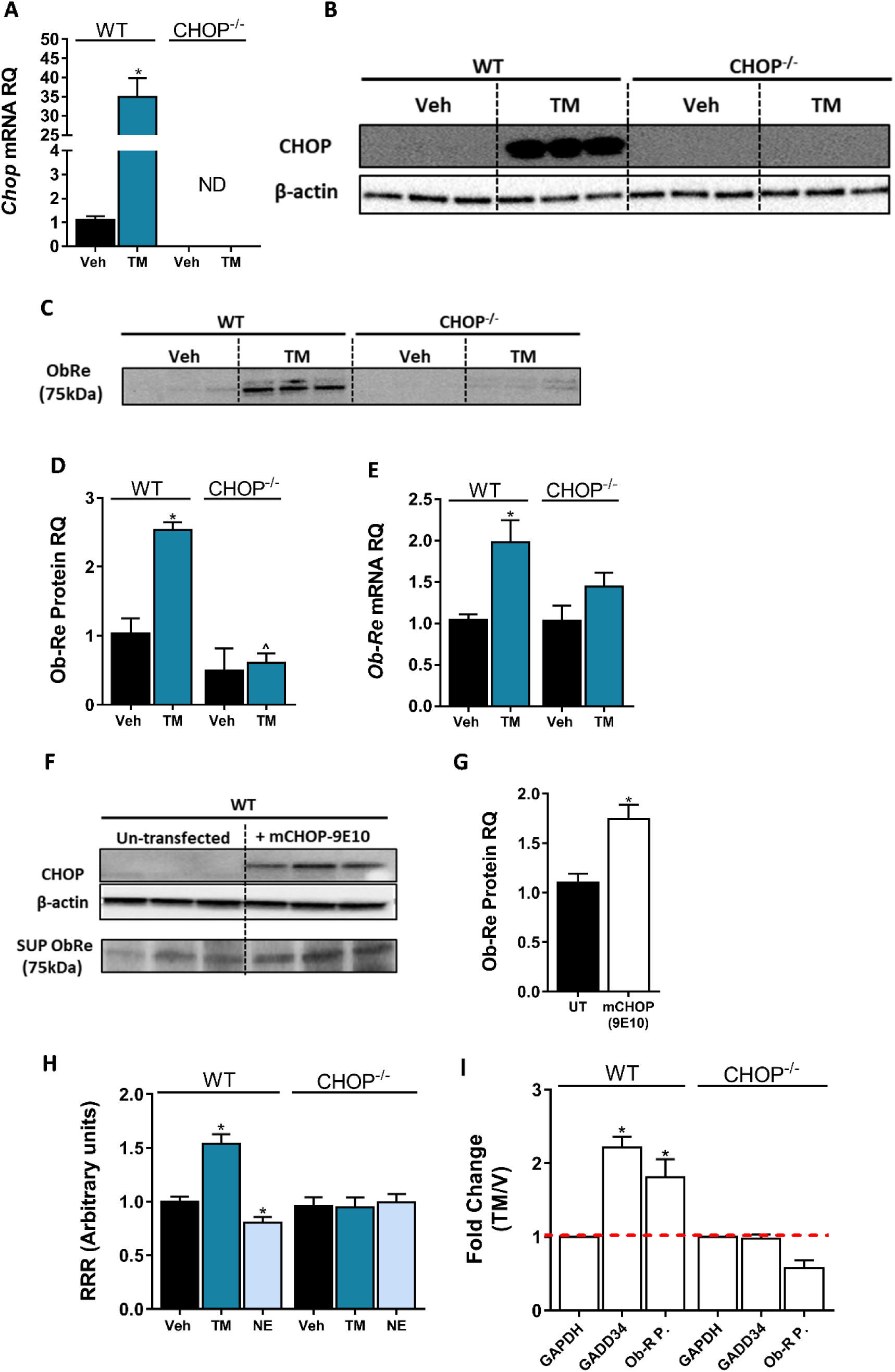
CHOP is a positive regulator of LEPR Promoter. Induction of CHOP mRNA **(A**, n=15) and protein **(B**, n=6) expression using a 6-hrs treatment with tunicamycin (TM; 2.5 μG/mL) was accompanied by elevated sOb-R levels in the culture media (blot was quantified using Ponceau staining as a loading control) **(C & D**, n=6) as well as mRNA levels **(E**, n=9-15) of WT hepatocytes. A transient CHOP overexpression induced elevation in sOb-R levels in the culture media of WT hepatocytes **(F & G**, n=3). Luciferase reporter assay **(H**, n=16) and Chromatin immunopecipitation (ChIP) **(I)** show increased LEPR promoter activity and CHOP binding to this promoter in WT hepatocytes treated with TM. Data represent mean ± SEM of indicated number of replicates in each panel from at least 2 independent experiments (and 3 independent ChIP assays). Blots are representative. *P<0.05 relative to vehicle treated cells. ^P<0.05 relative to the same treatment group of WT hepatocytes.

To further explore the possibility that CHOP can directly bind the LEPR promoter and control its expression, we preformed several ChIP experiments. *In silico* analysis of the LEPR promoter region revealed a putative binding site, corresponding to 5 of 6 nucleotides that compose a core sequence for CHOP binding (GRCm38:CM000997.2. Chromosome 4: 101,717,929-101,717,934) (Ubeda et al., 1996). As seen in Figure 6I, there was a 2-fold increase in the recovery of the qPCR product amplified with a primer set flunking the putative CHOP binding site, in cells that were treated with TM. A similar enrichment was seen in *Gadd34*, a well-known target of CHOP. This increase was limited to WT hepatocytes, validating the specificity of CHOP IP. Taken together, our data suggests that CHOP is able to occupy the LEPR promoter and directly regulate sOb-R levels in response to HFD consumption and/or CB_1_R activation.

The molecular signalling pathway(s) by which eCBs/CB_1_R regulates CHOP levels calls for further investigation. Nevertheless, one putative mechanism may involve Trib3, a multifunctional protein upregulated during ER stress by the PERK-ATF4-CHOP pathway, which mediates cell death. Trib3 represses its own expression by inhibiting the transcription of ATF4 and CHOP (Jousse et al., 2007; Mathur et al., 2014; Ohoka, Yoshii, Hattori, Onozaki, & Hayashi, 2005). In addition, many studies describe Trib3 as a key factor in mediating the anti-tumor effect of cannabinoids [reviewed in (Velasco, Sanchez, & Guzman, 2016)]. Our *in vivo* data indicate that HFD induces the mRNA and protein expression levels of hepatic Trib3, and that treatment with JD5037 restores these levels. This effect is limited to WT mice, while in CHOP KO mice, Trib3 levels did not change in response to HFD nor JD5037 treatment. Similarly, a direct activation of CB_1_R using NE upregulated Trib3 expression in WT, but not in CHOP KO hepatocytes (Supplementary Figure 3A-E), suggesting that Trib3 is indeed induced via CB_1_R signalling, and negatively regulates CHOP levels.

## Discussion

Since only free leptin crosses the blood-brain-barrier (BBB) and induces leptin signaling, the soluble isoform of leptin receptor, which sequesters free leptin in the serum and is considered as the main binding protein for leptin in the circulation, practically regulates leptin’s bioavailability and activity and can potentially affect leptin sensitivity/resistance. This is also true for peripheral tissues, where sOb-R/leptin complexes cannot bind to and activate membrane anchored leptin receptors. Many human and animal studies have demonstrated that sOb-R levels are inversely correlated with plasma levels of leptin, BMI and adiposity (Chan et al., 2002; Lahlou et al., 2000; Laimer et al., 2002; Ogier et al., 2002; Reinehr et al., 2005), suggesting that low levels of the soluble isoform contribute to obesity-related hyperleptinemia and subsequently, leptin resistance. In contrast to pathological conditions with a positive energy balance (i.e. obesity), human clinical situations associated with energy deficiency (i.e. starvation and/or anorexia nervosa) are characterized by upregulated circulating levels of sOb-R (Monteleone, Fabrazzo, Tortorella, Fuschino, & Maj, 2002; Reinehr et al., 2005; Stein, Vasquez-Garibay, Kratzsch, Romero-Velarde, & Jahreis, 2006; Zepf et al., 2012). Moreover, individuals carrying a mutated allele of LEPR, which leads to enhanced shedding of the leptin binding domain, have normoleptinemia and they are not obese (Lahlou et al., 2002). For these reasons, the sOb-R most likely plays a key role in the formation of central and peripheral leptin resistance conditions. Yet, only limited knowledge exists about the molecular mechanisms that regulate sOb-R production and secretion.

Using multiple cultured cell types, Gan and colleagues have shown that TNFα may induce cell surface expression of Ob-Rb as well as sOb-R levels (Gan et al., 2012). In addition, an *in vitro* study has demonstrated that increasing concentration of recombinant sOb-R diminishes STAT3 phosphorylation in response to leptin stimulation, but pre-incubation of leptin with recombinant sOb-R forms ligand-receptor complexes that do not affect leptin-mediated STAT3 phosphorylation (G. Yang et al., 2004). *In vivo*, it has been described that leptin stimulation as well as food deprivation specifically induce the expression of sOb-R in mouse liver (Cohen et al., 2005). It has been also demonstrated that in contrast to mice, the human sOb-R, exclusively generated through proteolytic cleavage of the extracellular domain of membrane-anchored isoforms (Maamra et al., 2001), is shed into the circulation by two well-known proteolytic enzymes, ADAM10 and ADAM 17, belong to the ‘ADAM’s family’ [review in (Schaab & Kratzsch, 2015)]. As we could not detect significant up/down regulation in the expression levels of these proteins in our experimental paradigms (Supplementary Figure 4), and the fact that we detected all changes in both mRNA and protein levels, we reasoned that the observed alterations in the circulating levels of sOb-R result from altered hepatic expression and secretion of the LEPR gene rather than decreased shedding. In addition, numerous stimuli, such as activation of protein kinase C, an increase in intracellular calcium, lipotoxicity and apoptosis, may contribute to the proteolytic cleavage of the extracellular leptin receptor domain (Maamra et al., 2001; Schaab et al., 2012). However, to our best knowledge, a molecular mechanism responsible for the decreased expression/shedding of sOb-R in obesity was never reported. Here, we describe, for the first time, the involvement of the eCB/CB_1_R system in regulating sOb-R levels and consequently leptin’s activity.

The importance of the eCB/CB_1_R system in regulating normal energy homeostasis as well as mediating obesity-related comorbidities is well acknowledged [review in (Simon & Cota, 2017)]. In fact, its pivotal interaction with leptin has been first described in 2001, demonstrating that leptin reduces the content of hypothalamic eCBs (Di Marzo et al., 2001), and attenuates eCB-mediated ‘retrograde’ neuronal CB_1_R signaling (Jo et al., 2005; Malcher-Lopes et al., 2006). On the other hand, activating CB_1_R by eCBs may, in turn, regulate leptin levels and signaling. Indeed, we have previously shown that CB_1_R activation in adipocytes and pre-junctional sympathetic fibers innervating the adipose tissue stimulates leptin biosynthesis and release, and its activation in the proximal tubules of the kidney inhibits leptin degradation and renal clearance (Tam et al., 2012), thus possibly contributing to leptin resistance. In agreement with these findings, peripheral CB_1_R blockade has been shown to ameliorate obesity-related hyperleptinemia, and subsequently restores leptin sensitivity in obese mice (Tam et al., 2012). By using both pharmacological and genetic approaches that target hepatic CB_1_R, our findings here suggest another novel mechanism by which the eCB system may regulate hepatic leptin resistance. Specifically, peripheral blockade as well as hepatic deletion/overexpression of CB_1_R modulate the expression levels of the sOB-R isoform in hepatocytes and its subsequent release into the circulation, reversing the CB_1_R-mediated decrease in sOb-R levels and hepatic leptin resistance during obesity. One should point out that although liver-specific CB_1_R KO mice retained higher levels of circulating sOB-R when fed a HFD, they were equally susceptible to DIO as their WT controls. Similarly, hepatic-specific CB_1_R transgenic mice in the CB_1_R-null background remained resistant to DIO while displayed significantly lower circulating sOB-R levels, as compared to their littermates. These data suggest that while liver CB_1_R expression is a major contributor to circulating sOB-R levels, their roles in regulating systemic/central energy balance will need to be further validated. Nevertheless, in accordance with our findings, Palomba and colleagues reported that CB_1_R activation interferes with leptin’s activity in hypothalamic ARC neurons (Palomba et al., 2015). On the other hand, opposite findings were reported by Bosier and colleagues, demonstrating that pharmacological or genetic deletion of CB_1_R in astrocytes downregulates Ob-Rb expression and leptin-mediated functional responses, whereas JZL195 (a dual MAGL and FAAH inhibitor) upregulates these features (Bosier et al., 2013). These differences can be explained by the distinct roles hepatocytes, astrocytes, and neurons play in peripheral and central metabolic regulations, and by cell-specific roles for CB_1_R in this regulation. As our findings demonstrate a similar effect of hepatic CB_1_R activation/overexpression or blockade/deletion on the different isoforms of LEPR, it seems equally possible that hepatic CB_1_R may affect DIO-related hepatic leptin resistance by not only modulating sOB-R levels, which controls leptin’s activity, but also by modulating the expression of Ob-Rb in hepatocytes. Further investigations would allow us to differentiate between these two pathways.

Obesity is often characterized by an ER stress and consequently an adaptive unfolded protein response (UPR), operated by three parallel sensors: activating transcription factor 6 (ATF6), inositol requiring enzyme 1α (IRE1α), and protein kinase R-like ER kinase (PERK) (Walter & Ron, 2011). The activation of the latter induces the phosphorylation of eIF2α, which, in turn, inhibits transcription and protein synthesis (Ron, 2002). In case of an extreme ER stress conditions, CHOP is activated by the PERK signaling pathway, and executes ER stress-mediated apoptosis (Hu et al., 2018; Zinszner et al., 1998). In fact, ER stress has been shown to contribute to the development of hypothalamic leptin resistance, by impairing the transport of leptin across the BBB and suppressing STAT3 phosphorylation (El-Haschimi, Pierroz, Hileman, Bjorbaek, & Flier, 2000; Hosoi et al., 2008; L. Ozcan et al., 2009; X. Zhang et al., 2008). In addition, under physiological conditions, excess nutrients increases the demand for protein synthesis by the liver, leading to ER stress and UPR activation, which resolves the stress within hours (Oyadomari, Harding, Zhang, Oyadomari, & Ron, 2008). Nevertheless, chronic ER stress in the liver was demonstrated in both obese mice and human (U. Ozcan et al., 2004; Puri et al., 2008). Here, we demonstrate that obese mice had elevated levels of phosphorylated eIF2α, indicating increased ER stress; an effect that was reversed by peripheral CB_1_R blockade and was absent in LCB1^-/-^. These findings are in agreement with our previous reports, where we reported that a neutral CB_1_R antagonist (AM6545) has the ability to reduce the HFD-induced upregulation in hepatic eIF2α (Tam et al., 2010), and that hepatic activation of CB_1_R induces ER stress and contributes to insulin resistance (Liu et al., 2012). Unexpectedly, we found that the hepatic gene and protein levels of CHOP were significantly decreased in obese mice, and were upregulated by peripheral CB_1_R blockade. This observation is conceptually in agreement with several previous reports that describe a modulated UPR signaling with altered sensitivity or output that might implicate conditions of persistence/repeated stress (Chambers, Petrova, Tomba, Vendruscolo, & Ron, 2012; Gomez & Rutkowski, 2016; Preissler et al., 2015; L. Yang et al., 2015).

Apart from its role in ER stress-mediated apoptosis, CHOP has been implicated in regulating other processes such as inflammation (Endo, Mori, Akira, & Gotoh, 2006; Nakayama et al., 2010), insulin resistance (Maris et al., 2012; Song, Scheuner, Ron, Pennathur, & Kaufman, 2008) and adiposity. Specifically in the liver, Chikka and colleagues suggest that CHOP is a suppressor of key regulators of lipid metabolism like *Cebpα Pparα* and *Srebf1* (Chikka, McCabe, Tyra, & Rutkowski, 2013), and demonstrate that CHOP deficient mice tend to develop hepatic steatosis in response to bortezomib-induced ER stress. This is in agreement with an earlier report describing higher body weight and adiposity in female CHOP KO mice compare to WT controls (Ariyama et al., 2007). In contrast, we show that male CHOP KO mice and their WT littermate controls gain comparable amount of weight and have similar body composition following exposure to an HFD for 14 weeks. Nevertheless, we did see a trend toward increased liver triglycerides.

An interesting observation was that the eCB ‘tone’ of lean and obese CHOP KO mice was comparable. To our best knowledge, a direct regulation of eCB synthesis or degradation by CHOP has never been described. Thus, our data imply a possible link between the two. Interestingly, JD5037 failed to reverse many of the metabolic abnormalities, such as HDL and LDL content as well as liver triglycerides in DIO CHOP KO mice. It also had much smaller effect on total body fat mass then in WT DIO mice. Whereas basal circulating levels of sOb-R were comparable between WT and CHOP KO mice, its levels were markedly lower in the liver of CHOP KO mice as well as cultured hepatocytes. Moreover, sOb-R remained low in these mice even on HFD, supporting a role for CHOP in regulating the synthesis of sOb-R in the liver. The failure of JD5037 to elevate sOb-R levels in obese CHOP KO mice places CHOP downstream of CB_1_R in this molecular cascade.

As mention earlier, the molecular signalling pathway(s) by which eCBs or CB_1_R regulates CHOP levels is outside the scope of this work. However, two possible mechanisms might be relevant. The first putative mechanism may involve Trib3. Interestingly, it has been shown that Δ^9^-THC as well as synthetic cannabinoid agonists upregulate Trib3 expression (Blazquez et al., 2006; Carracedo et al., 2006; Salazar et al., 2013; Vara et al., 2011) to engage apoptosis in variable cancer models. Moreover, Cinar *et al*. demonstrated that hepatic CB_1_R induces ER stress in hepatocytes by increasing *de novo* synthesis of ceramides (Cinar et al., 2014), which are also involved in Trib3 upregulation following ER stress (Carracedo et al., 2006). In line with our observation that Trib3 levels are negatively correlated with sObR and are elevated following direct or indirect activation of CB_1_R, we may suggest Trib3 to be the molecular linker between the two pathways. Second, CHOP is a cAMP responsive protein, so its expression is induced via a cAMP response element (CRE) (Conkright et al., 2003; Pomerance et al., 2003; Ramji & Foka, 2002; Wilson & Roesler, 2002). CB_1_R, a G-protein coupled receptor (GPCR), which upon activation recruits Gi protein, can inhibit the activity of adenylyl cyclase and reduce the levels of cAMP (Turu & Hunyady, 2010). It is therefore plausible that a decline in cAMP following CB_1_R activation inhibits CHOP transcription. This hypothesis is more appealing if one considers the pivotal role of cAMP in regulating liver metabolism (Wahlang, McClain, Barve, & Gobejishvili, 2019), and takes into account the fact that reduced levels of cAMP were documented in HFD-fed mice (Zingg et al., 2017). Yet, further studies will need to explore the specific molecular pathways linking together hepatic eCB/CB_1_R system and CHOP.

In conclusion, we report a new role for the hepatic eCB/CB_1_R in the development of hepatic leptin resistance, by reducing the expression and/or subsequent release of sOb-R (Figure 7). We show that peripherally restricted CB_1_R antagonism has the ability to restore sOb-R levels, contributing to the reversal of obesity-induced hyperleptinemia. We also suggest that upon CB_1_R blockade in hepatocytes, CHOP levels are upregulated via reduced Trib3 expression. CHOP, in turn, directly binds the LEPR promoter and promotes the expression of sOb-R.

**Figure 7.**
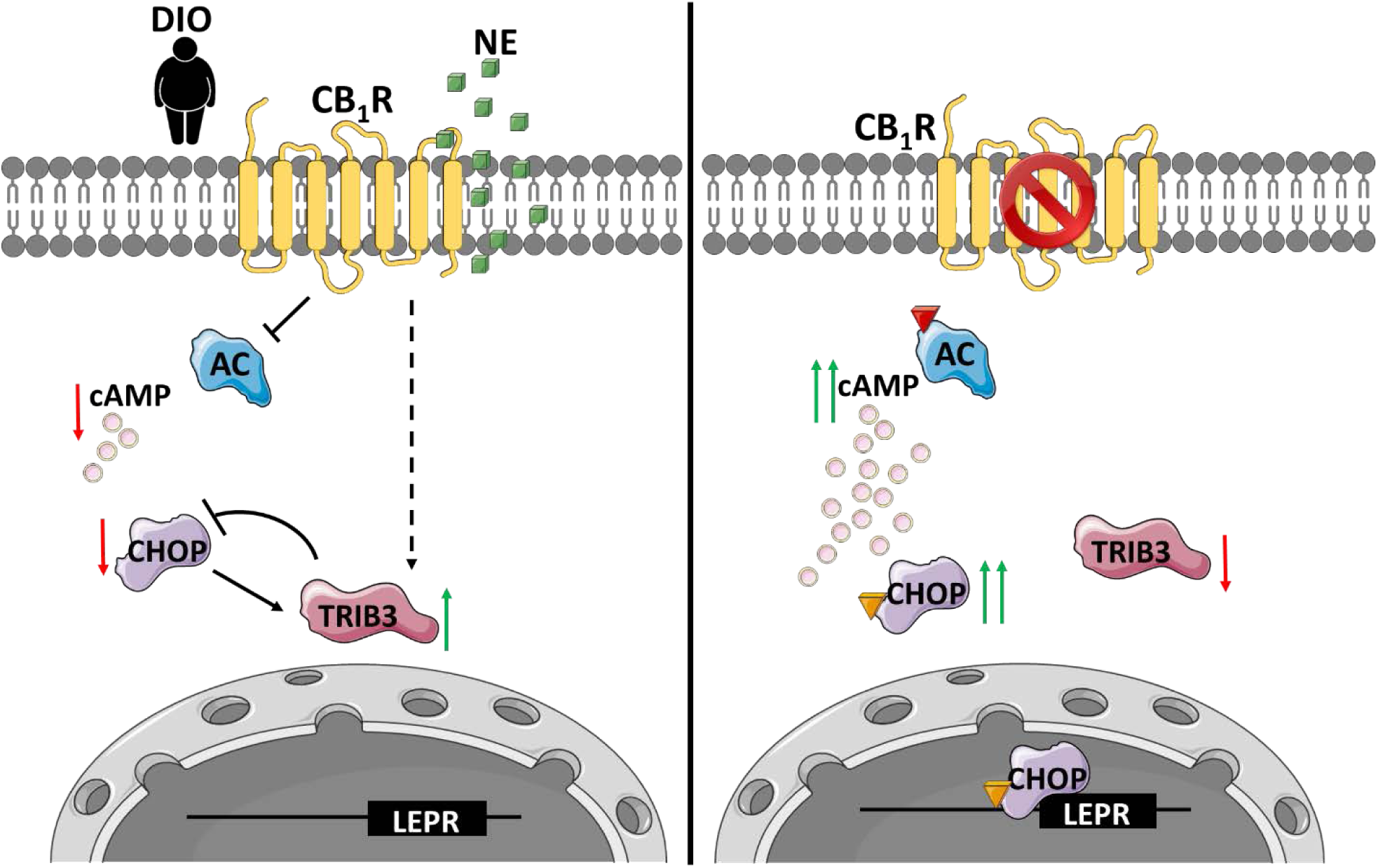
An illustration that describes the suggested molecular mechanism involving CB_1_R and CHOP in the regulation of sOb-R levels. **(Left)** When activated by eCBs upregulated during diet-induced obesity (DIO) or synthetic cannabinoids, such as noladin ether (NE, green squares), CB_1_R attenuates cAMP (pink circles) production by inhibiting adenylate cyclase (AC), as well as upregulates Trib3 expression. As a consequence, CHOP levels are reduced, transcribing less LEPR. **(Right)** Blocking CB_1_R in hepatocytes reverses these changes, leading to the activation and translocation of CHOP to the nucleus, which, in turn, directly binds the LEPR promoter and promotes the expression of sOb-R. Red arrows represent downregulation, whereas green arrows represent upregulation. Coloured triangles represent activation.

## Materials and Methods

### Animals and Experimental Protocol

All animal studies were approved by the Institutional Animal Care and Use Committee of the Hebrew University of Jerusalem (AAALAC accreditation #1285). C57Bl/6 (Envigo, Israel), LCB1^-/-^ and hTgCB1^-/-^ (kindly provided by Dr. George Kunos, NIH) or B6.129S(Cg)-Ddit3^tm2.1Dron^/J (CHOP KO, The Jackson Laboratory #005530), and their WT littermate controls were used for *in vivo* experiments. All mice were male, 8-10-week-old at the beginning of each experiment. To generate DIO (body weight > 42g), mice were fed with a standard diet (STD; 14% fat, 24% protein, 62% carbohydrates; NIH-31 rodent diet) or a high-fat diet (HFD; 60% fat, 20% protein and 20% carbohydrates; Research Diet, D12492) for 14 weeks. Treatment with JD5037 (3 mg/kg, ip) or vehicle (1% Tween80, 4% DMSO, 95% Saline) was conducted for 7 days, and 12 hrs following the last dose, the mice were euthanized by a cervical dislocation under anesthesia, and blood and livers were harvested for further analyses. For leptin sensitivity test, mice were fasted for 24 hrs before an ip administration of recombinant mouse leptin (3 mg/kg). 1 hr following leptin administration, mice were euthanized, liver were harvested and processed for phosphorylated STAT3 detection using Western blot.

### Cell Culture

WT or CHOP KO immortalized hepatocytes [described in (Uzi et al., 2013)] were maintained in DMEM (01-100-1A; Biological Industries, Israel) supplemented with 5% FCS, 100 mM Glutamine, 100 mM Na-Pyruvate, and Pen/Strep. Cells were cultured at 37°C in a humidified atmosphere of 5% CO_2_/95% air. To test the effect of CB_1_R activation, cells were seeded in 6-well plates (25×10^4^ cells/well) for 24 hrs. Then, growth medium was replaced with a serum-free medium (SFM) for an additional 12 hrs. At the morning of the experiment the medium was replaced with fresh SFM containing either vehicle (EtOH), 2.5 μM noladin ether (NE; Cayman Chemicals, Ann Arbor, Michigan) or a combination of 100 nM JD5037 (Haoyuan Chemexpress Co., Ltd) and 2.5 μM NE. 24 hrs later, cells were harvested for further analyses as described below.

### Measurements of sOb-R

Soluble leptin receptor was quantified by an ELISA kit, capable to differentiate the soluble isoform from other isoforms, according to manufacturer’s instructions (E03S0226; Shanghai Bluegene Biotech, China). Briefly, for serum measurements, we diluted serum in saline (1:2) and 100 μL from the diluted samples were analyzed. For hepatic measurements, 50-100 mg tissue samples were homogenized in 300 μL of 1xPBS and centrifuged for 5 min in 5000 rpm. 100 μL of cleared lysates were analyzed. Data were normalized to sample protein content, determined using the Pierce™ BCA Protein Assay Kit (Thermo Scientific, IL).

To measure sOb-R protein content in cell culture media, we used trichloroacetic acid (TCA) precipitation protocol. 350 μL of 100% TCA were added to 1.6 mL culture media, vortexed and incubated for 30 min on ice. Samples were then centrifuged to pellet proteins (14000 rpm, 10 min, 4°C). Pellets were washed in 100% acetone, resuspended in 0.1 M NaOH and protein loading dye, and analyzed by Western blot. Ponceau staining of the blots was used as loading control for quantification.

### Validation of LEPR antibody

The specificity of the anti-LEPR antibody was validated in a control experiment (Supplementary Figure 5), where mouse LEPR was overexpressed in kidney cell line by using a viral infection. The viral vector encoded Ad-GFP-mLEPR (ADV-263380, VECTOR BIOSYSTEMS Inc.) was used in a multiplicity of infection of 50, and cells were harvested for Western blot analysis 24 hrs post infection.

### Real-time PCR

For total mRNA isolation, tissue samples or hepatocytes were washed in 1xPBS and harvested using Bio-Tri RNA lysis buffer (Bio-Lab, Israel). Extracted RNA was treated with DNase I (Thermo Scientific, IL), and reverse transcribed using the Iscript cDNA kit (Bio-Rad Laboratories, CA). Quantitative PCR reactions for *ObRe*, *Chop* or *Gadd34* were performed using iTaq Universal SYBR Green Supermix (Bio-Rad Laboratories, CA), and the CFX connect ST system (Bio-Rad Laboratories, CA). Relative quantity (RQ) values of all tested genes were normalized to *Ubc*. Primers are listed in Supplementary Table 1.

### Western blot Analysis

Tissue samples or hepatocytes were washed in cold 1xPBS, and harvested in a RIPA buffer (25 mM Tris-HCl pH 7.6, 150 mM NaCl, 1% NP-40, 1% sodium deoxycholate, 0.1% SDS), vortexed and incubated for 30 min at 4°C, then centrifuged for 10 min at 14,000 rpm. Protein concentrations were determined using the Pierce™ BCA Protein Assay Kit (Thermo Scientific, IL). Cleared lysates were supplemented with protein sample buffer, resolved by SDS-PAGE (4-15% acrylamide, 150V) and transferred to PVDF membranes using the Trans-Blot® Turbo™ Transfer System (Bio-Rad Laboratories, CA). Membranes were incubated for 1 hr in 5% milk (in TBS-T) to block unspecific binding, washed briefly, and incubated overnight at 4°C with the following primary antibodies: LEPR (NB-120-5593, Novus), phosphorylated STAT3 (9145, Cell Signaling), STAT3 (9139, Cell Signaling), phosphorylated eIF2α (9721, Cell Signaling), eIF2α (9722, Cell Signaling), CHOP (2895S, Cell Signaling), Trib3 (ab137526, Abcam) or β-Actin (ab49900, Abcam). Anti-rabbit (ab97085, Abcam) or mouse (ab98799, Abcam) horseradish peroxidase (HRP)-conjugated secondary antibodies were used for 1 hr at room temperature, followed by chemiluminescence detection using Clarity™ Western ECL Blotting Substrate (Bio-Rad Laboratories, CA). Densitometry was quantified using ImageLab software. Protein RQ was calculated as the ratio between LEPR to total protein signal (ponceau) in culture media supernatants or to β-Actin in cell and tissue lysates.

### Body composition

Total body fat and lean masses were determined by EchoMRI-100H™ (Echo Medical Systems LLC, Houston, TX, USA).

***HDL and LDL measurements*** were done using the Cobas C-111 chemistry analyzer (Roche, Switzerland).

### Hepatic Triglycerides Measurements

Tissue lipids were extracted as described in (Folch, Lees, & Sloane Stanley, 1957), and quantified using Triglyceride Assay Kit (ab65336; Abcam). Data were normalized to tissue weight.

### Endocannabinoid measurements by LC-MS/MS

eCBs were extracted, purified, and quantified in liver homogenates, as described previously (Drori et al., 2019; Udi et al., 2017). LC-MS/MS was analyzed on an AB Sciex (Framingham, MA, USA) Triple Quad™ 5500 mass spectrometer coupled with a Shimadzu (Kyoto, Japan) UHPLC System. eCBs were detected in a positive ion mode using electron spray ionization (ESI) and the multiple reaction monitoring (MRM) mode of acquisition. The levels of each compound were analyzed by monitoring multiple reactions. The molecular ion and fragment for each compound were measured as follows: m/z 348.3→62.1 (quantifier) and 91.1 (qualifier) for AEA, m/z 379.3→287.3 (quantifier) and 91.1 (qualifier) for 2-AG. The levels of AEA and 2-AG in samples were measured against standard curves, and normalized to tissue weight.

### Chop Overexpression

WT hepatocytes were transfected with mCHOP-WT-9E10-pCDNA1 vector (Addgene plasmid #21913) using Lipofectamin 3000. Cells were harvested 24 hrs post-transfection and CHOP expression was validated by Western blot analysis.

### Luciferase Promoter Assay

Mus musculus Ob-R promoter sequence (GRCm38:CM000997.2. Chromosome 4: 101,716,750-101,718,250 forward strand) was cloned into pGL3-basic vector (E1751, Promega). Reporter vector, as well as Renilla luciferase vector, were then co-transfected into WT or CHOP KO hepatocytes using Lipophectamin 3000. 24 hrs post transfection, cells were treated with either vehicle, 2.5 μg/mL tunicamycin (11089-65-9; Holland Moran, Israel) or 2.5 μM NE for indicated period. At the end of the experiment, luciferase activity was measured using Dual-Glo® Luciferase Assay System (E2920, Promega). Data are presented as the ratio between Firefly and Renilla luciferase activity.

### Chromatin Immunoprecipitation (ChIP)

WT or CHOP KO hepatocytes were seeded in 100 mm plate and left to adhere (5×10^6^ cells per plate; 3 plates for each sample). The next day, cells were treated with either DMSO or 2.5 μg/mL tunicamycin for 6 hrs to induce CHOP expression. At the end of incubation, cells were washed in 1xPBS, fixed with 1% formaldehyde for 10 min, then quenched with 125 mM glycine and harvested from plates. Following centrifugation, cells were re-suspended in a lysis buffer and sonicated for 12 cycles of 30 sec pulse followed by 30 sec rest in 70% amplitude. Sheared DNA was diluted in a ChIP dilution buffer and pre-cleared with magnetic ProteinG-sepharose beads for 4 hrs at 4°C. 10% of the lysate was removed and saved as “Input”. The rest of the lysate was divided and each part was incubated overnight at 4°C with 2.5 μg of either anti-H3 (ab1791, Abcam), anti-CHOP (2895S, Cell Signaling) or IgG isotype control. Antibody-chromatin complexes were precipitated with magnetic protein G-Sepharose beads, washed with low salt, high salt, lithium chloride and Tris-EDTA buffers. DNA was then eluted from beads, digested with proteinase K and purified. 1.5 μL of clean DNA was used in a qPCR reaction using specific primers for GAPDH or LEPR promoter region. For a positive control, GADD34 primers were used. For each sample, we calculated the ratio between the RQ (expressed as % of input) of ObR promoter qPCR product in αCHOP IP relative to αH3 IP. This ratio was normalized to the ratio of GAPDH qPCR product to control for non-specific binding. Data are expressed as the fold-change of this ratio in tunicamycin-treated compare to vehicle-treated cells. ChIP primers are listed in Supplementary Table 2.

### Statistics

Data are presented as mean ± SEM. Unpaired two-tailed Student’s t-test was used to determine variations between groups. Results in multiple groups were compared by ANOVA followed by a Bonferroni test (GraphPadPrism v6 for Windows). Significance was set at P < 0.05.

## Acknowledgments

This study was supported by grants from the Israeli Science Foundation (ISF; 617/14 and 158/18) and The Obesity Society’s Early Career Research Award to J.T.

## Author Contributions

AD conducted the experiments and analyzed the data. AG, SA, LH and DW assisted with animal care and *in vivo* experiments. RH and GS provided reagents and technical assistance. AN conducted the LC-MS/MS analysis. MS provided essential knowledge and reagents for ChIP analysis. BT provided CHOP KO mice and cultured hepatocytes. AD and JT designed and supervised the experiments and wrote the manuscript.

## Conflict of Interest

The authors have declared that no conflict of interest exists.

## Inventory of Supplemental Material

The following Supplemental Figures and Tables provide additional information supporting the role of hepatic CB_1_R in regulating sOb-R levels via CHOP:

**Supplementary Figure 1.**
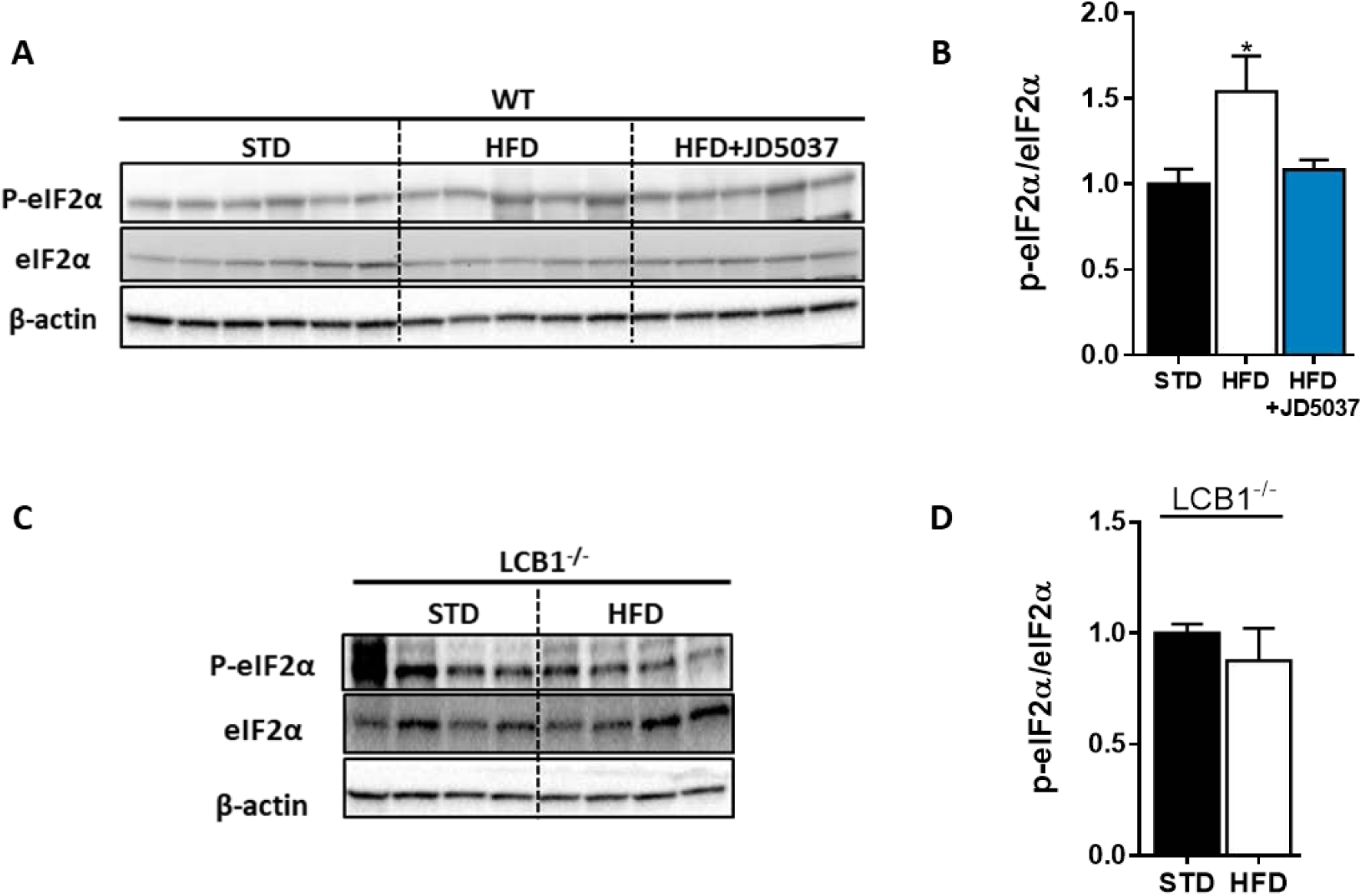
HFD induces eIF2α phosphorylation in WT mice. Western blot analysis **(A, C)** and quantification **(B, D)** shows increased hepatic ratio between phosphorylated and total eIF2α in WT, but not LCB1^-/-^ mice. Data represent mean ± SEM of 5-6 samples in each group. *P<0.05 relative to STD fed animals from the same strain. Blots are representative.

**Supplementary Figure 2.**
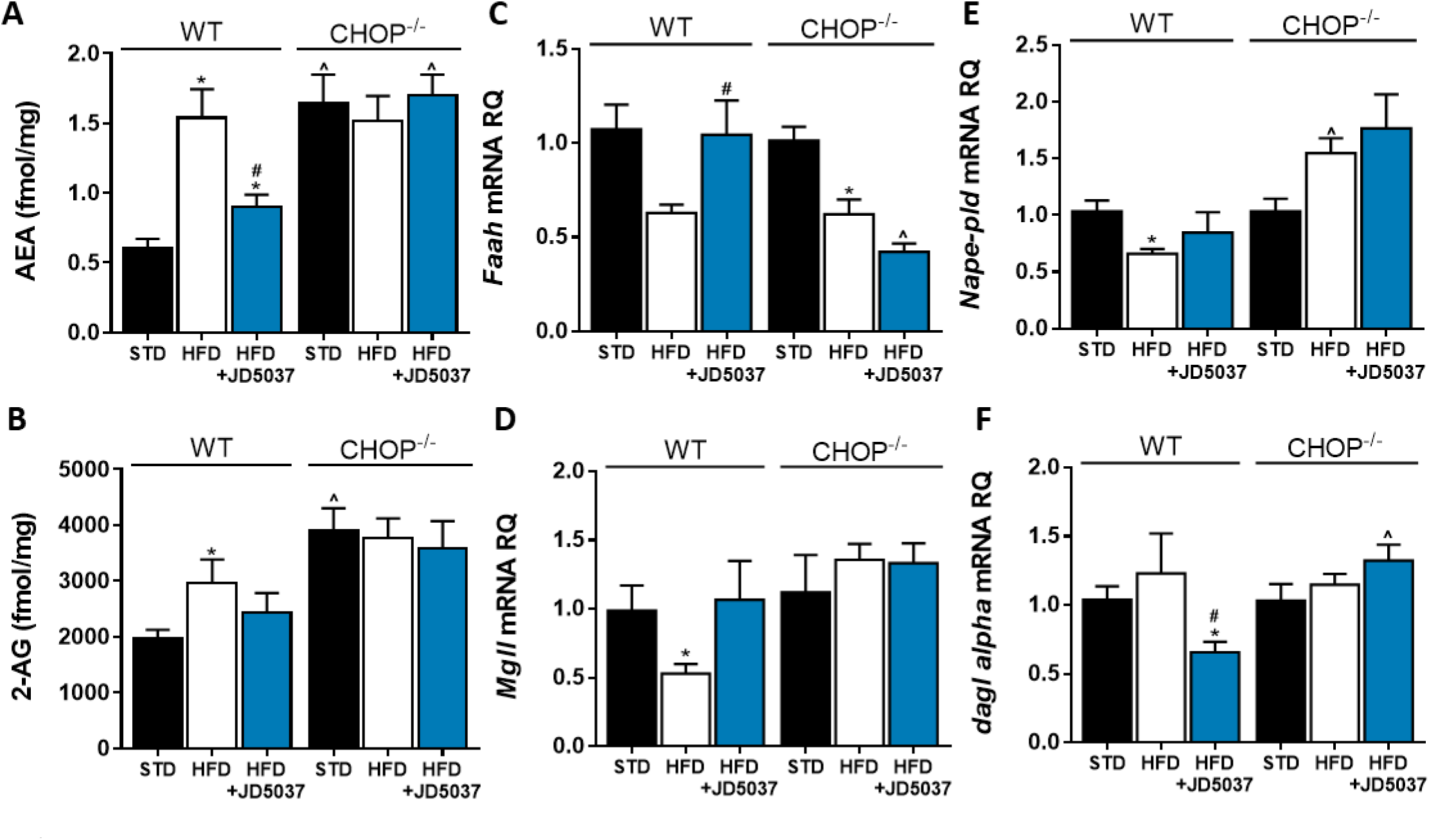
An altered endocannabinoid ‘tone’ in CHOP deficient mice. Measurements of hepatic AEA **(A**, n=6-12) and 2-AG **(B**, n=6-12) levels in mice demonstrate an expected elevation of eCB levels in obese mice, and decreased levels following JD5037 treatment (3 mg/kg, for 7 days) in WT, but not CHOP KO mice. This observation is partly explained by the changes measured in the mRNA expression levels of their synthesis and degradating enzymes **(C-F**, n=6-15). Data represents mean ± SEM of indicated number of replicates in each panel. *P<0.05 relative to STD fed animals from the same strain. #P<0.05 relative to HFD fed mice from the same strain. ^P<0.05 relative to the same treatment group of WT mice.

**Supplementary Figure 3.**
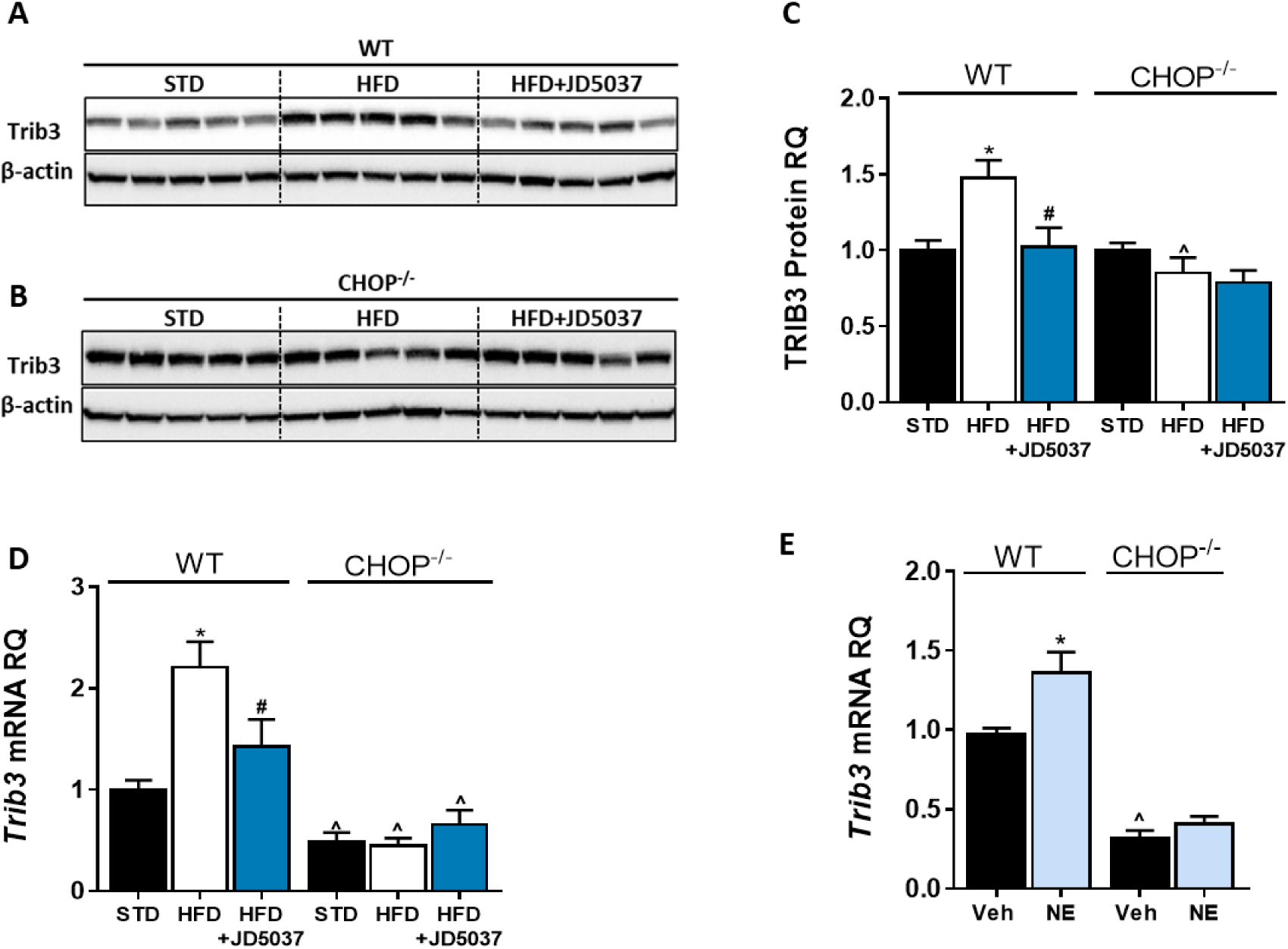
Trib3 as a possible link between CB_1_R and CHOP. Western blot analysis and quantification **(A-C**, n=5) and mRNA of hepatic Trib3 expression levels **(D**, n=5) in WT and CHOP KO mice show increased expression in obese WT, but not LCB1^-/-^ mice. A direct activation of CB_1_R by using noladin ether (NE) upregulated the mRNA expression of Trib3 in WT, but not in CHOP KO hepatocytes. *In vivo*: Data represent mean ± SEM of indicated number of replicates in each panel. Western blots are representative. *P<0.05 relative to STD fed animals from the same strain. #P<0.05 relative to HFD fed mice from the same strain. ^P<0.05 relative to the same treatment group of WT mice. *In vitro:* Data represent mean ± SEM of 12-15 replicates from at least 3 independent experiments. *P<0.05 relative to vehicle treated cells in the same genotype. ^P<0.05 relative to same treatment paradigm in WT.

**Supplementary Figure 4.**
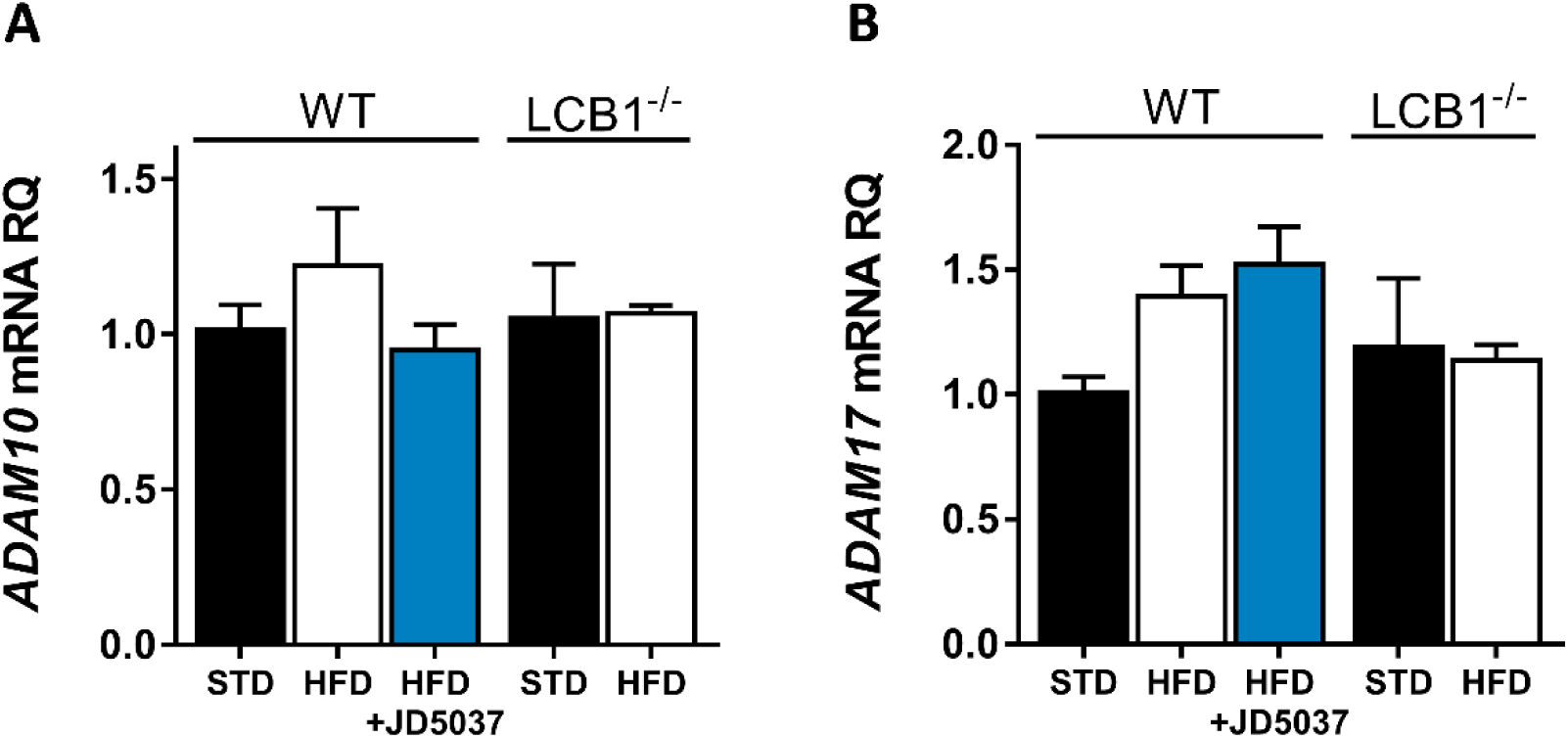
Hepatic expression of ADAM10 and ADAM17. Hepatic mRNA expression levels of ADAM10 **(A)** and ADAM17 **(B)** in WT mice fed with STD or HFD for 14 weeks and treated with JD5037 (3 mg/kg, for 7 days) as well as in LCB1^-/-^ mice fed either STD or HFD for the same period. Data represent mean ± SEM of 5-6 animals per group.

**Supplementary Figure 4.**
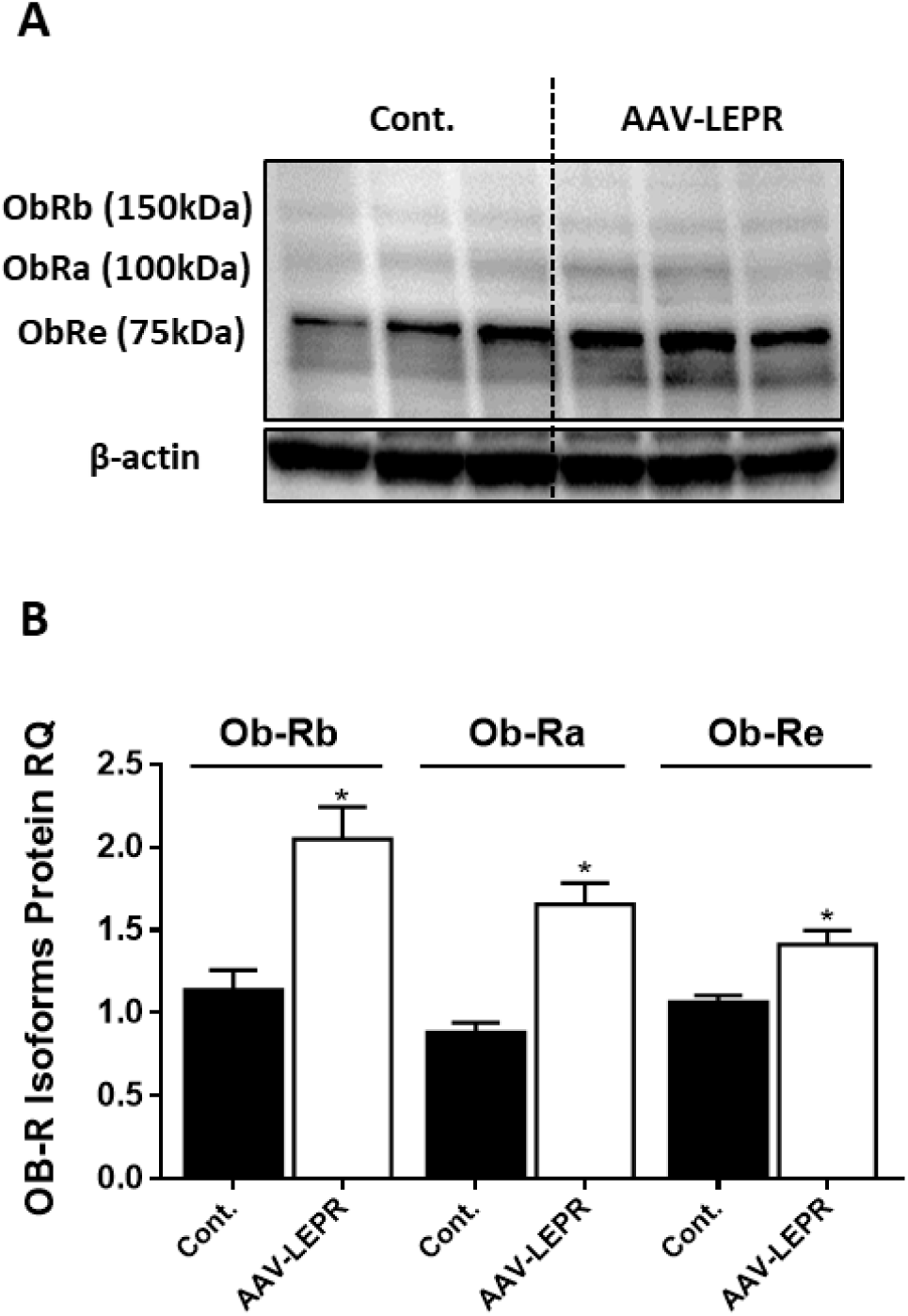
LEPR antibody detects isoforms of 75, 100 and 150 kDa. Western blot analysis and quantification **(A**-**B**, n=3) of LEPR expression levels in HK-2 cell line, infected with either empty vector or mouse LEPR expressing AAV. Vector also expressed eGFP and infection efficiency was approximately 30 percent, as assessed by fluorescent microscopy (not shown). Data represent mean ± SEM of indicated number of replicates in each panel. *P<0.05 relative to control cells.

**Supplementary Table 1.**
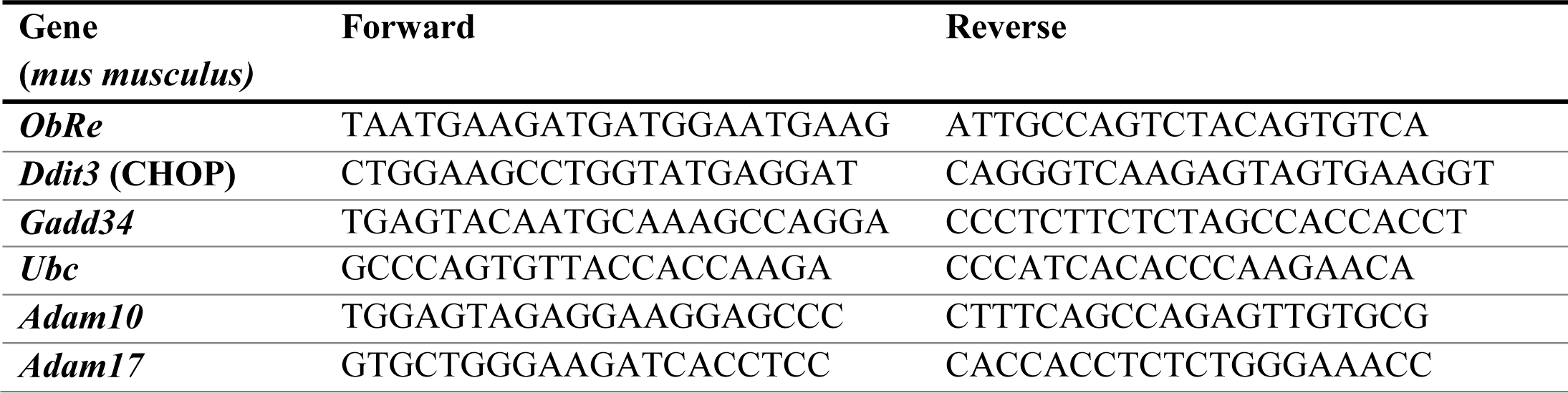
mRNA Primers

**Supplementary Table 2.**
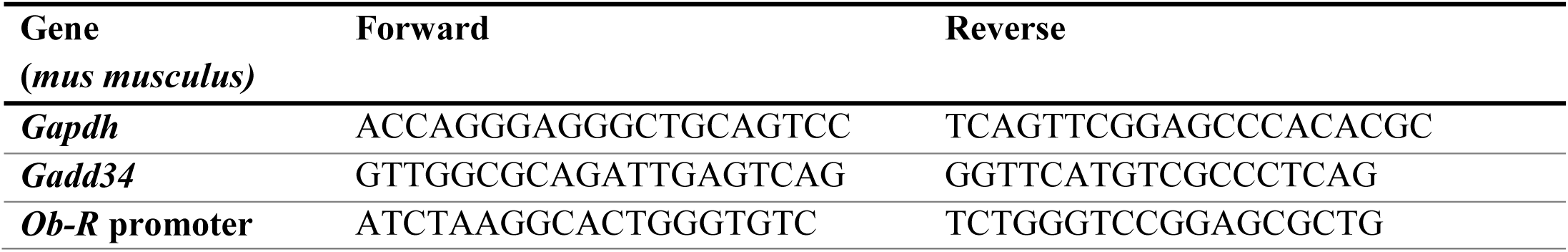
ChIP primers

## Notes

### Competing Interest Statement

The authors have declared no competing interest.

